# The Rad52 superfamily as seen by AlphaFold

**DOI:** 10.1101/2024.08.09.607149

**Authors:** Ali Al-Fatlawi, Md. Ballal Hossen, Stella de Paula Lopes, A. Francis Stewart, Michael Schroeder

## Abstract

Rad52, a highly conserved eukaryotic protein, plays a crucial role in DNA repair, especially in double-strand break repair. Recent findings reveal that its distinct structural features, including a characteristic *β*-sheet and *β*-hairpin motif, are shared with the lambda phage single-strand annealing proteins, Red*β*, indicating a common superfamily. Our analysis of over 10,000 single-strand annealing proteins (SSAPs) across all kingdoms of life supports this hypothesis, confirming their possession of the characteristic motif despite variations in size and composition. We found that archaea, representing only 1% of the studied proteins, exhibit most of these variations. Through the examination of four representative archaeal SSAPs, we elucidate the structural relationship between eukaryotic and bacterial SSAPs, highlighting differences in *β*-sheet size and *β*-hairpin complexity. Furthermore, we identify an archaeal SSAP with a structure nearly identical to the human variant and screen over 100 million unannotated proteins for potential SSAP candidates. Our computational analysis complements existing sequence with structural evidence supporting the suggested orthology among five SSAP families across all kingdoms: Rad52, Red*β*, RecT, Erf, and Sak3.

## 2 Introduction

Rad52 is a nearly ubiquitous eukaryotic protein involved in DNA repair, particularly the repair of double-strand breaks by facilitating the pairing of complementary DNA strands [1, 2]. It is involved in both homologous recombination (HR) and single-strand annealing (SSA) pathways and has been extensively studied in vitro for its abilities to form undecameric rings in the absence of DNA [3, 4] and to promote annealing of complementary DNA strands. [5]. These properties are also presented by the lambda phage single-strand annealing protein (SSAP), Red*β* [6]. Although other phage SSAPs form filaments rather than rings [7], Passy et al. speculated that Rad52 and Red*β* are functionally related. Contrary to assumptions, Kharlamova et al. found that RAD52 facilitates single-strand annealing and homology detection primarily through short oligomers rather than ring structures. SSAPs were classified into three main classes named after their leading members, namely Rad52, Red*β*/RecT, and Erf [8] as well as Sak3. Using advanced bioinformatic tools, Erler et al. [9] identified a distant tripartite similarity between Rad52 and Red*β* that coincided with the most conserved sequences within the Rad52 class, suggesting an orthologous relationship between these two very distantly separated SSAPs. Several other observations also enhanced the suggested orthology amongst these SSAPs: (i) the single-strand DNA binding/annealing domains of these SSAPs occupy the N-terminal 180 amino acids, which includes the tripartite amino acid signature; (ii) in the absence of DNA, they multimerize into rings or chains at high concentrations in vitro (*>*0.5 M); (iii) they bind ssDNA with modest affinity but have only low affinity for double-strand (ds) DNA; (iv) beyond the N-terminal annealing domain, the C-terminal regions are not required for annealing but are required for homologous recombination (HR) [10, 11]. Independently, the distant sequence relationship between Rad52 and Red*β*/RecT classes was confirmed using different bioinformatic methodologies [12], and the Erf class was included in the suggested SSAP superfamily [13]. The Rad52 superfamily hypothesis was recently strengthened by the cryo-EM structural resolution of two members of the Red*β*/RecT class [14, 15] that identified structural similarities with the known RAD52 structure [3, 4] and by AlphaFold predictions that led to the identification of a new protein fold [16]. This protein fold, which is shared by the Erf class, involves a well-conserved three-stranded *β*-sheet traversed on the inside by a helix. The outside of the *β*-sheet sets the curvature of the helical filaments and rings formed by these SSAPs, whereas the traversing helix is secured by a second helix with accompanying *β*-hairpin and another helix. These latter two secondary elements show considerable variability amongst various SSAPs. Al-Fatlawi et al. visually depicted the similarities and variations of this arrangement of five secondary structural elements across eight selected SSAPs [16].

Here, we refine the above approach and apply it across bacterial, archaea, and eukaryotic SSAPs. Following Illergard et al. [17], who argue that structure is three to ten times more conserved than sequence, we compare these SSAPs by their structure instead of their sequence. Such a large-scale comparison is only made possible by the recent large-scale availability of high-quality predicted protein structures in the AlphaFold database [18].

We will shed light on the overall phylogenies computed from sequence and from structure focusing in particular on the preservation of the central *β*-hairpin and *β*-sheet motif across all families. This is of particular interest in the light of SSAP distribution across superkingdoms: The majority of all known SSAPs are bacterial, complemented by eukaryotic Rad52s and a small number of archaea sequences. We quantify variation in the core SSAP structural motifs across these distant relationships of bacteria, eukaryota, and archaea. Furthermore, we exploit the structural perspective to screen for potential novel SSAPs in the large body of millions of unannotated proteins.

## 3 Results

### 10,280 SSAPs predicted structures

Our study presents a comprehensive structural analysis of the Rad52 single-strand annealing protein (SSAP) superfamily and provides insights into their structural diversity. We retrieved the members of single-strand annealing protein families Rad59/52/22 (Rad52 for short throughout the entire manuscript), Erf, Sak3, Red*β*, and RecT from the Interpro database [19] and enriched them with 3D structures predicted by AlphaFold [20], [21]. After filtering out low-quality structures (see methods), we focused on 10,280 high-quality predicted protein structures with 5,150 Red*β* and RecT family members and a roughly equal number from Rad52, Erf, and Sak3.

### Most SSPs are bacterial

Next, we enriched the data with phylogenetic information. An interesting hypothesis on the origin of eukaryotic Rad52 can be drawn from the very imbalanced breakdown by superkingdoms (see Table 1): 90% of the total SSAPs surveyed are of bacterial origin. The remaining 10% are shared by eukaryotes (9%) and archaea (1%). While the 1% archaea structures cover the whole five families, the eukaryotic SSAPs are predominantly limited to Rad52. This is interesting since Rad52 is also present in bacteria and in archaea. Our data suggest a scenario where the full diversity of the SSAP families possibly originated in bacteria and archaea first, with eukaryotic Rad52 evolving from ancestral forms found in both bacterial and archaea lineages. Subsequent analysis will explore how these evolutionary relationships manifest themselves in the structural characteristics of SSAPs.

**Table 1:** Distribution of Rad52, RecT, Redß, Erf, and Sak3 Families Across Archaea, Eukaryota, and Bacteria.

| **Family** | **Archaea** | **Eukaryota** | **Bacteria** | **Total** |
| --- | --- | --- | --- | --- |
| Rad52 | 13 | 854 | 1,178 | 2045 |
| RecT | 25 | 5 | 3,860 | 3890 |
| Redβ | 8 | 0 | 1,252 | 1260 |
| Erf | 63 | 2 | 2,582 | 2647 |
| Sak3 | 15 | 0 | 423 | 438 |
| **Total** | **124** | **861** | **9,295** | **10,280** |

### Structure and sequence phylogenetic trees largely agree

Protein families within the InterPro database are delineated based on sequence data, utilizing sophisticated algorithms such as hidden Markov models. We conducted comparisons across 10,280 single-strand annealing proteins (SSAPs) using both sequence alignment via the Blast algorithm and structural alignment via the state-of-the-art TM-align algorithm to ascertain the consistency between structural and sequence-based classifications. The dendrograms generated from sequence and structure comparisons are shown in Fig. S1. Both clusterings clearly separate all five families and in particular Rad52 and Red*β*. The clustering patterns consistently differentiated eukaryotic from bacterial Rad52. However, discrepancies arose in the placement of a Rad52 subgroup termed RDM, attributed to its possession of an additional RNA binding motif. Despite this subgroup’s outlier status, our findings underscore the overall agreement between structural and sequence-based classifications of SSAPs.

### Archaea SSAPs are representative of all SSAPs

Among the single-strand annealing proteins (SSAPs) analyzed, less than 1% were attributed to the archaea superkingdom, known for harboring extremophiles. It is still an open question on how eukaryotes evolved from a world with only two domains, bacteria and archaea [22]. Given the evolutionary significance of archaea in understanding eukaryotic origins, we sought to discern potential structural distinctions between archaeal, bacterial, and eukaryotic SSAPs. The question arises whether the few archaea SSAP structures differ significantly from bacterial and eukaryotic ones or not. Thus, we clustered all SSAPs by their structure and highlighted archaea SSAPs (see Figure 1A). The 124 archaea SSAPs spread evenly across the full tree of 10280 structures and are thus representative for the whole data. Since one focus of our analysis is the relation of bacterial and eukaryotic SSAPs, we decided to use archaea structures as independent reference points. Utilizing archaeal structures as independent reference points, we further distilled this set to 12 representative structures through clustering. Manual inspection and refinement led to the selection of four representative structures, each showcasing variations in *β*-sheet and *β*-hairpin motifs. These motifs exhibited diverse characteristics, including variations in strand length and complexity of the hairpin structure. Two representatives belonged to the Rad52 family, while one each represented RecT and Red*β*, respectively. These underscore the structural diversity present within archaeal SSAPs.

**Fig. 1:**
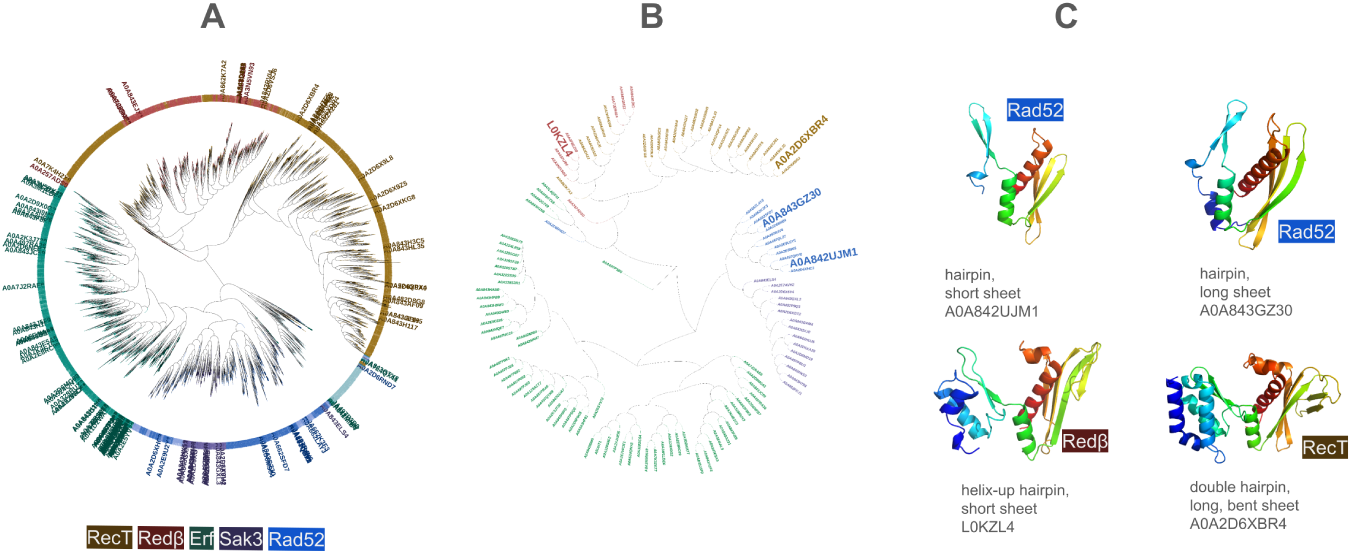
(A) 10280 SSAPs clustered by structural similarity confirm the definition of SSAP families RecT (brown), Red*β* (red), Erf (green), Sak3 (purple), and Rad52 (blue). RecT/Red*β* are clearly separated from Erf, Sak3, and Rad52. Sak3 is mixed within Rad52. Eukaryotic (lightblue) and prokaryotic (dark blue) Rad52 are mixed and clearly separated from the Rad52 subfamily RDM1 (turquoise), which has an additional RNA-binding motif. Out of the 10280 SSAPs depicted, the 124 archaea SSAPs are labeled by their IDs, which shows that the 124 covers the full diversity of the 10280. (B) Close-up of the 124 archaea SSAPs clustered by structural similarity with the four selected representatives labeled by their identifier. (C) The four selected representatives with their *β*-hairpin and *β*-sheet motif. They will serve as references in the subsequent analyses.

### Four representative SSAPs align with experimental templates

For the Rad52, RecT, and Red*β* families, which have experimentally determined 3D protein structures from human and two bacterial phages, we investigated the extent to which the evolutionarily distant archaea representatives align with them. Using the TM-score, a measure of structural similarity ranging from 0 to 1 (values *>* 0.5 indicating similarity), we obtained promising scores: 0.51 for Rad52, 0.65 for RecT, and 0.55 for Red*β*. In stark contrast, sequence identities on aligned motifs were notably low, ranging from 5 to 8%. These findings underscore a remarkable phenomenon: despite minimal sequence similarity, the structural resemblance among these proteins is striking.

These results refer to the monomeric structures, but SSAPs oligomerize and form regular, circular quaternary structures. Rad52 e.g. forms a ring of 11 monomers [23]. This begs the question whether the predicted archaea Rad52 representatives are consistent with such a quaternary ring structure? This could be expected since the ring structure is essential for the function in single strand annealing, but it would also be surprising since neural networks for 3D structure prediction such as AlphaFold only train on monomers and not on oligomers. To answer this question, we used the experimentally determined oligomeric Rad52 structure with its 11 monomers as a template and superimposed 11 copies of the predicted representative archaea Rad52 structures. To assess whether the hypothetical ring structure of predicted Rad52 is realistic we counted atom clashes between neighboring monomers. We found that for the long *β*-sheet Rad52 representative only 9 out of 869 atoms clash at a distance cut-off of 2Å. For the short *β*-sheet representative it is equally low at 8 clashes out of 749 atoms. We proceeded similarly for Red*β* and RecT. Experimental structures for the latter two SSAPs form large helical structures instead of rings [24], [25]. We found that the predicted archaea Red*β* representative leads to 16 out of 1087 atom clashes when superposed to the Red*β* template and the RecT representative to 12 out 1236 for the RecT template. This is a very remarkable result as it suggests that AlphaFold correctly predicts aspects of quaternary structure without having seen quaternary structures. Conceptually this would imply that primary structure not only codes for tertiary structure, but also quaternary structure.

### Do Redß and RecT form a single group as suggested by Iyer et al.?

Iyer et al. conducted sequence analyses of single-strand annealing proteins (SSAPs), proposing that RecT and Red*β* constitute a single family distinct from the other three families. Our sequence and structure analyses confirm the distinction between RecT and Red*β*, although they exhibit closer structural similarity to each other than to the other families (see Figure 1A and Fig. S1A and 1B). To understand these relations in detail, we plotted the structural similarity of all SSAPs against the archaea RecT and Red*β* representatives (see Figure 2). Structural similarity ranges from 0 (not similar) to 1 (identical) and a value of 0.5 indicates a similar fold [26], [27]. Nearly all of the RecT and Red*β* SSAPs have a structural similarity of 0.5 and better to both of the two RecT and Red*β* representatives. In contrast, the other three families, Rad52, Erf, Sak3, are all below 0.5 similarity to either of the two RecT and Red*β* representatives. These findings are in agreement with Iyer et al. However, all of the Red*β* SSAPs are more similar to the Red*β* than to the RecT representative and nearly all RecT SSAPs are more similar to the RecT than to the Red*β* representative. This clear separation supports the notion that RecT and Red*β* should be treated as distinct families, contrary to the proposal by Iyer et al.

**Fig. 2:**
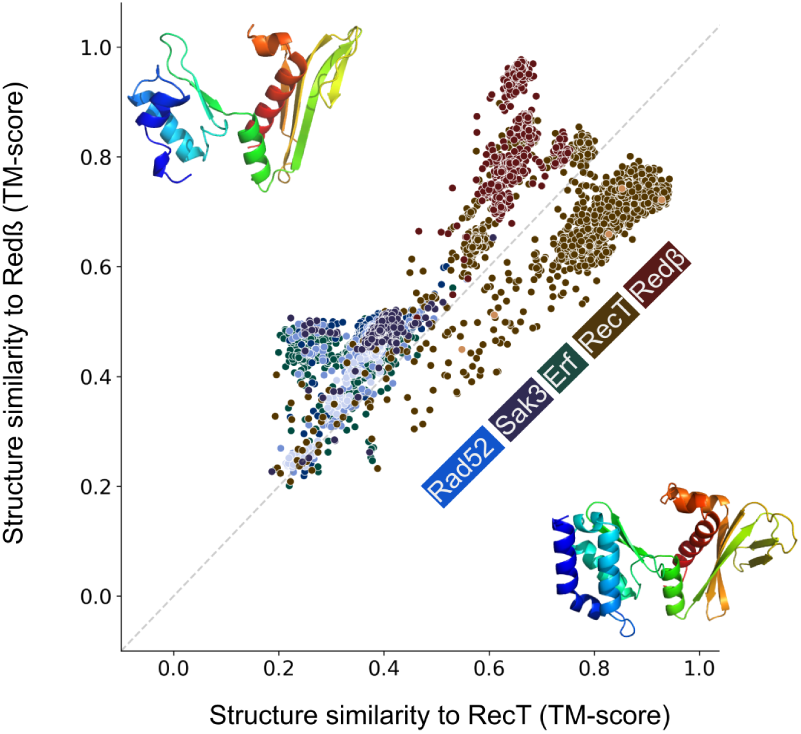
Scatter plot of structural similarity (TM-score) of each of the 10280 SSAP against RecT (brown) and the Red*β* (red) representatives. RecT and Red*β* SSAPs are clearly separate from the rest but also clearly separate from each other.

### How do Red**β** and RecT compare to Rad52 and in particular to eukaryotic Rad52?

Iyer et al. pointed out that RecT/Red*β* are clearly distinct from the other three families. This is also supported by the clusterings in Figure 1A and Fig. S1A and B. To examine the details, we compared structural similarity of all SSAPs against the remaining two representative archaea structures, which both belong to the Rad52 family and which differ in the lengths of the three *β*-strands in the characteristic *β*-sheet motif (see Figure 3). In contrast to the previous analysis, all but some RecT structures share a structural similarity of 0.5 or better with both representatives and can hence be considered similar. As expected, Rad52, Erf, and Sak3 have higher similarities to the representatives. Interestingly, nearly all of the Red*β*, RecT, and Sak3 SSAPs are more similar to the short *β*-sheet than to the long *β*-sheet representative. In contrast, Rad52 spreads along two diagonal axes representing similarity to long vs. short *β*-sheets on the one hand and to similarity to either of those two vs. none of these two. Regarding the axis short vs. long *β*-sheets, bacterial Rad52 breaks down into 376 SSAPs (35%), which are more similar to the long *β*-sheet representative and 802 SSAPs (65%), which are more similar to the short *β*-sheet representative. Eukaryotic Rad52 is mostly more similar to the long *β*-sheet representative. The second axis of spread of Rad52 is characterized by poor similarity to either of the two representatives. In fact, the Rad52 SSAPs concerned form a Rad52 subfamily called RDM1, which contains an additional RNA recognition motif. Poor similarity against this motif explains the spread. Taken together, these results suggest that the Ur-Rad52 is of bacterial origin with short *β*-sheets.

**Fig. 3:**
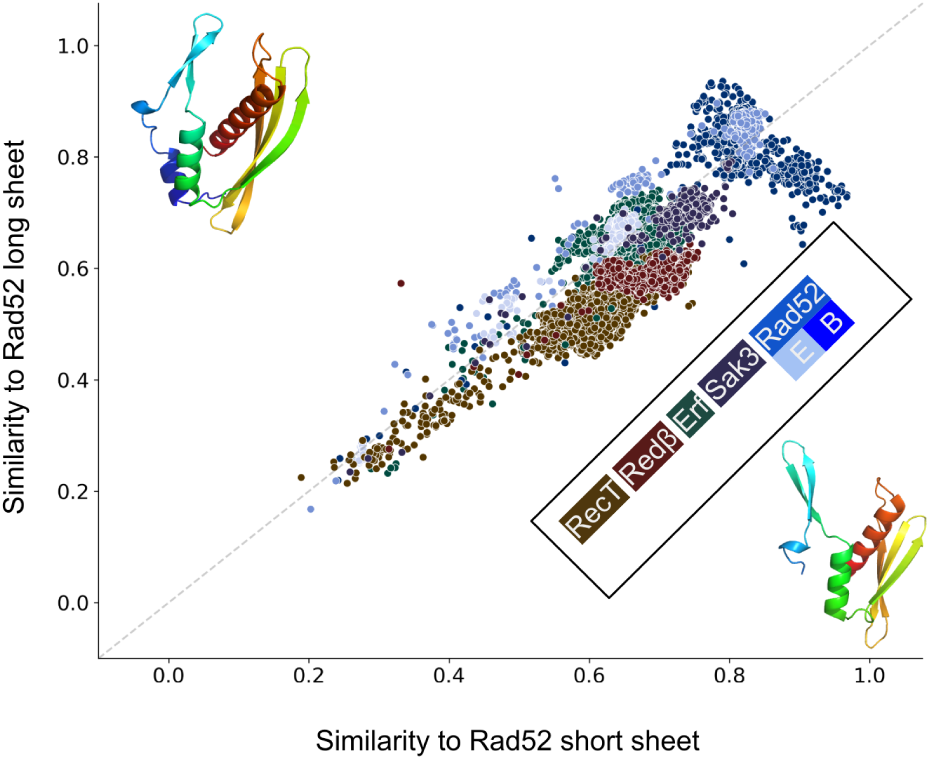
Scatter plot of structural similarity (TM-score) of each of the 10280 SSAP against the long and short *β*-sheet Rad52 representatives. Most SSAPs show a good similarity. Bacterial Rad52 SSAPs (dark blue) are highly similar to both long and short *β*-sheet Rad52, while eukaryotic Rad52 (medium blue) is also highly similar to both but more similar to long *β*-sheet Rad52. The Rad52 subfamily RDM1 (light blue) is due to its additional RNA-binding domain more dissimilar to both.

### Do all SSAPs exhibit the characteristic motif?

Single-strand annealing proteins are defined by their function. This function is closely linked to the *β*-hairpin and the *β*-sheet motif, both of which play a role in DNA binding. Do all of the 10280 SSAPs contain these two motifs? As can be seen from Figure 1C, they are present in the four selected representatives.

Out of the 10,280 SSAPs analyzed, 208 have a TM-score below 0.5, indicating dissimilarity in structure. Another 2,995 SSAPs fall within the TM-score range of 0.5 to 0.7, suggesting moderate structural similarity. Remarkably, the majority, comprising 7,077 SSAPs, demonstrate a TM-score above 0.7, indicating high structural similarity among this significant portion of the analyzed SSAP dataset (see Supplementary Table S1). Visual inspection confirmed that a TM-score of greater than 0.5 implies that the *β*-sheet motif is very well conserved and that the *β*-hairpin motif is present. A score of greater than 0.7 indicate that both motifs are well conserved. Overall, we found that out of the 10280 SSAPs 98% have a TM-score of greater than 0.5 to at least one of the four representatives (see Table 2). For 69% it is even greater than 0.7. For 50% of the SSAPs the TM-score is greater than 0.5 against at least three of the four representatives. From this we conclude that all SSAPs contain the two characteristic motifs and that it is constituting for single-strand annealing. This comprehensive examination underscores the fundamental role of the *β*-hairpin and *β*-sheet motifs in the function of SSAPs, highlighting their significance in single-strand annealing processes.

**Table 2:** Number of SSAPs with good TM-score against 1, 2, 3, or 4 representatives.

|  | 10,280 SSAPs |  |  | Unannotated Proteins |  |  |
| --- | --- | --- | --- | --- | --- | --- |
|  | >0.5 | >0.6 | >0.7 | >0.5 | >0.6 | >0.7 |
| >0 | 10,072 | 9,947 | 7,077 | 206,859 | 8,136 | 3,143 |
| >1 | 10,003 | 9,218 | 3,876 | 61,842 | 5,937 | 1,797 |
| >2 | 5,146 | 3,011 | 3 | 8,148 | 763 | 0 |
| >3 | 3,663 | 473 | 0 | 2,131 | 99 | 0 |

### Quantifying motif alignment lengths across all families

To quantify the structural diversity and conservation of the *β*-hairpin and *β*-sheet motifs across the single-strand annealing protein families, we employed two approaches. Firstly, we analyzed the distribution of alignment lengths against the representatives. Secondly, we quantified the composition of helix, strand, and loop residues in the motifs. Fig.Figure 1C visually illustrates the variation in size and shape of the *β*-hairpin and *β*-sheet motifs using four representatives from archaea. In Figure 4, we present the alignment length of each of the 10,280 SSAPs grouped by family, with each of these representatives depicted separately. Overall, alignments range from length 60 to 180. The most pronounced solitary peak can be found for the RecT representatives and the RecT family members (Figure 4A). Their alignments are nearly all around 170 residues long. This confirms that the RecT family is structurally coherent and distinct from Red*β*. Figure 4B also shows that the next longest pronounced peak is Red*β* at an alignment length of 120 to 140. All of the other bacterial SSAPs match the RecT representative with lengths of 80 to 100 residues and only eukaryotic Rad52 stands out with alignments of only 60 residues. When considering Red*β* as representative (Figure 4B) a consistent picture emerges, in which RecT/Red*β* are clearly separated from the rest. The bulk of RecT and Red*β* family members match the Red*β* representative with a length of around 130, but there is a smaller group of Red*β* matching their representatives better with a larger length of 145. Interestingly, eukaryotic Rad52 fits better to the Red*β* representative than to the RecT representative with alignments peaking at length 80 instead of 60. Turning to the Rad52 representatives (Figure 4C and D), with the longer and shorter *β*-sheet motif, there is a remarkable difference. For the short *β*-sheet motif (4D), eukaryotic and bacterial Rad52 as well as the large Erf family match with an alignment length of over 100 in a very compact manner. The agreement of Erf and Rad52 also supports our choice of representatives. Although our method did not select any Erf family members as representatives, they are nonetheless structurally very well represented by the selected short *β*-sheet Rad52 representative. Turning to 4C, the long *β*-sheet representative matches for all families including RecT at a reduced alignment length to 90. Overall, this analysis supports that RecT and Red*β* are distinct from the other families, but can also be clearly separated from each other. Eukaryotic Rad52 can be distinguished from bacterial Rad52, which breaks down into two groups. One similar to eukaryotic Rad52 and one different. However, variations emerged in helix residues, with Sak3 notably low at 3, and Red*β* and RecT notably high at 28 and 42, respectively. This underscores the distinction between RecT/Red*β* and the others, as well as the difference between these two families. A similar trend was observed in the helix residues of the *β*-sheet motif, reflecting perpendicular helix length. Rad52, Erf, and Sak3 exhibited shorter helices (15 to 20) compared to Red*β* and RecT (26 and 32). Strand lengths also diverged, with eukaryotic Rad52 having 37 residues compared to bacterial Rad52’s 27. RecT and Red*β* approached eukaryotic Rad52’s strand length, while Erf and Sak3 were more aligned with bacterial Rad52. These findings illuminate how sequence alterations drive structural variation while preserving motif topology.

**Fig. 4:**
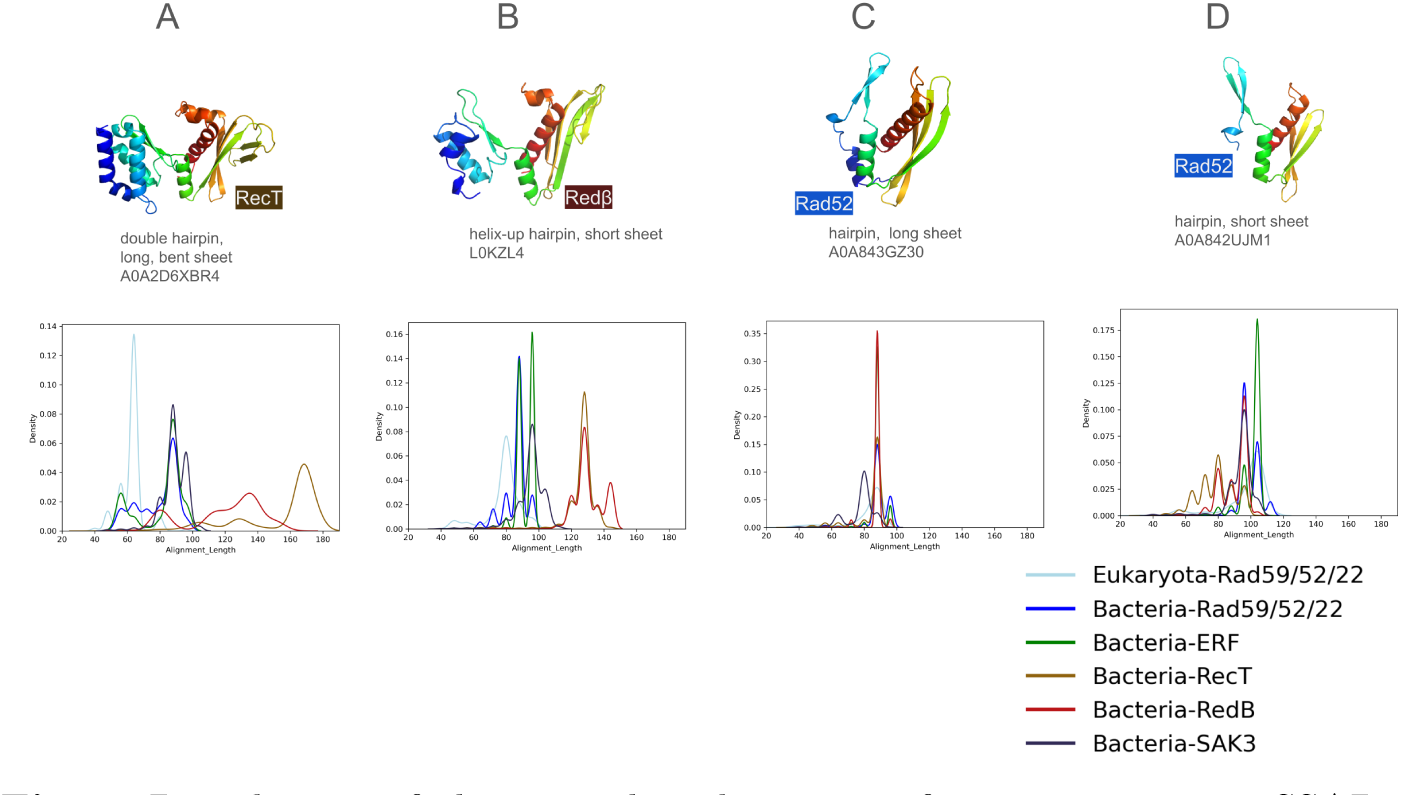
Distribution of alignment lengths against four representative SSAPs. Lengths of structural alignment against the four representatives. A) RecT B) Red*β*-C) Rad52 (long *β*-sheet) D) Rad52 (short *β*-sheet).

### Quantifying secondary structure composition of the motifs across all families

In further quantifying motif variation, we focused on secondary structure composition to highlight differences in helix and strand length among families. We limited the analysis to those families and superkingdoms with more than 100 members, i.e. all of the bacterial SSAPs and eukaryotic Rad52. Across families, the amount of strand residues in the *β*-hairpin motif does not vary strongly neither across families nor between eukaryotic and bacterial. It remaines relatively consistent, ranging from 11 to 15, (see Table 3). However, variations emerged in helix residues, with Sak3 notably low at 3, and Red*β* and RecT notably high at 28 and 42, respectively.This underscores the distinction between RecT/Red*β* and the others, as well as the difference between these two families.A similar trend was observed in the helix residues of the *β*-sheet motif, reflecting perpendicular helix length. Rad52, Erf, and Sak3 exhibited shorter helices (15 to 20) compared to Red*β* and RecT (26 and 32).Strand lengths also diverged, with eukaryotic Rad52 having 37 strand residues, i.e. around 12 per strand, compared to bacterial Rad52 27, i.e. 9 per strand. RecT and Red*β* approached eukaryotic Rad52’s strand length, while Erf and Sak3 were more aligned with bacterial Rad52. Taken together, these numbers document in detail how changes in sequence lead to variation in structure leaving the overall motif topology intact.

**Table 3:** Number of helix and strand residues in motif for eukaryotic Rad52 and bacterial Rad52, Red*β*, RecT, Erf, and Sak3.

| SS | SSAP | Superking. | β-sheet motif | | β-hairpin motif | |
| --- | --- | --- | --- | --- | --- | --- |
|  |  |  | median | std | median | std |
| H | RecT | Bacteria | 32 | 6.26 | 42 | 7.44 |
| H | Redβ | Bacteria | 26 | 3.57 | 28 | 6.95 |
| H | Erf | Bacteria | 20 | 3.03 | 15 | 4.80 |
| H | Rad52 | Bacteria | 17 | 2.87 | 12 | 3.76 |
| H | Rad52 | Eukaryota | 15 | 4.50 | 18 | 7.42 |
| H | Sak3 | Bacteria | 15 | 3.20 | 3 | 6.43 |
| S | Rad52 | Eukaryota | 37 | 8.06 | 11 | 10.32 |
| S | RecT | Bacteria | 35 | 8.18 | 12 | 5.25 |
| S | Redβ | Bacteria | 32 | 6.54 | 13 | 4.63 |
| S | Erf | Bacteria | 29 | 7.83 | 12 | 10.64 |
| S | Rad52 | Bacteria | 27 | 7.69 | 14 | 5.66 |
| S | Sak3 | Bacteria | 26 | 10.76 | 16 | 7.11 |

### Are there novel SSAPs?

The SSAPs documented in the Interpro database were assigned based on sequence information [19], [28]. Since structural information is more conserved than sequence [29], we expected there to be novel candidate SSAPs, which could not be identified by sequence-based methods. Therefore, we compared all 117,501,756 million proteins, which are listed in UniProt but not annotated by the Interpro database, against the four representatives. We found a total of 206859 proteins to have some structural similarity (TM-score greater than 0.5) to at least one of the four representatives. This number shrinks to 3143 at a more rigid cut-off of 0.7. The majority are highly similar to long (2164) and the short (1458) *β*-sheet Rad52. A minority of 667 and 655 proteins are structurally similar to RecT and Red*β* representatives, respectively (see FigFigure 5).

**Fig. 5:**
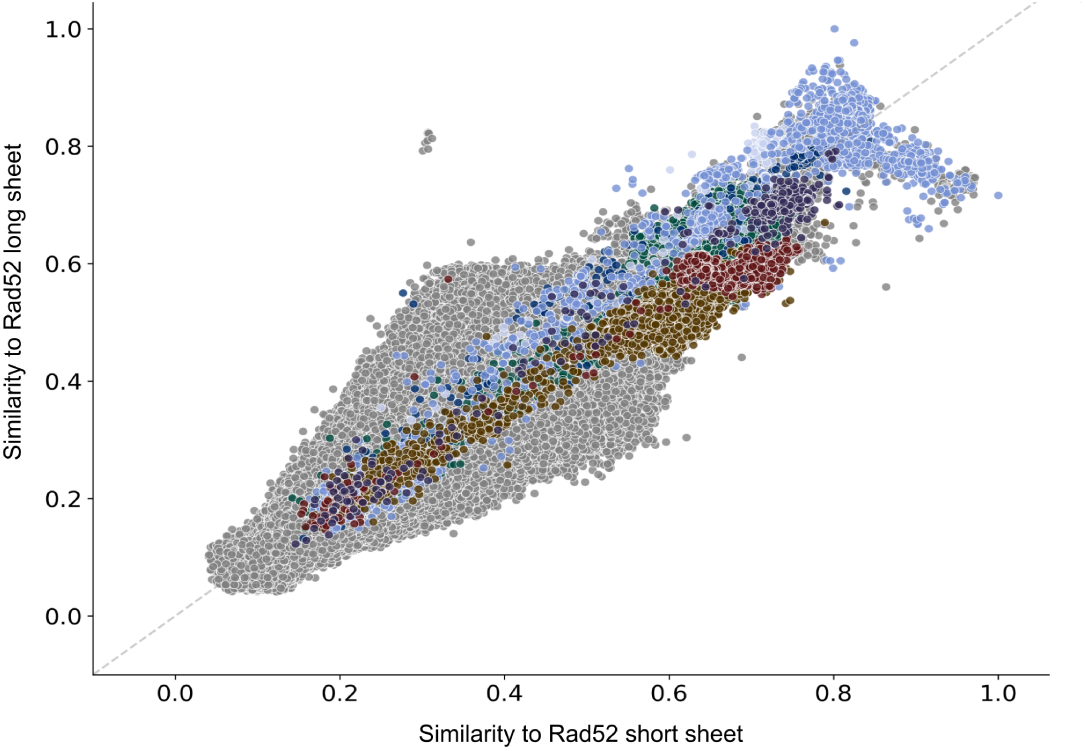
Scatterplot of structural similarity (TM-score) of all Alphafold structures including 117,5 million unreviewed ones against the two representative Rad52 SSAPs. The bulk of the proteins are below 0.5. Many of the known SSAPs have good scores better than 0.5. Among the top-scoring proteins, there are 2164 (1458) proteins with a TM-score greater than 0.7 against the long (short) *β*-sheet Rad52 representative. These are novel candidate SSAPs.

### Oceanic Archaeal SSAP Resembles Human Rad52

Remarkably, we identified an oceanic archaeal SSAP, UniProt ID A0A2D6XHC3, structurally resembling human Rad52 despite a low amino acid sequence identity. The oceanic archaeal SSAP, obtained from the Candidatus Pacearchaeota archaeon, was identified in a meta-genomic study of the Tara Oceans circumnavigation expedition [30]. Despite a sequence identity of only 30.63% at a coverage of 47%, effectively amounting to less than 15% sequence identity, the archaea and human structures exhibit remarkable structural similarity with a TM-score of 0.82 (see Figure 6). This finding underscores the remarkable conservation of structural features across evolutionary distant organisms and suggests functional conservation in their roles as single-strand annealing proteins. It highlights the importance of considering structural information alongside sequence data when studying protein evolution and function.

**Fig. 6:**
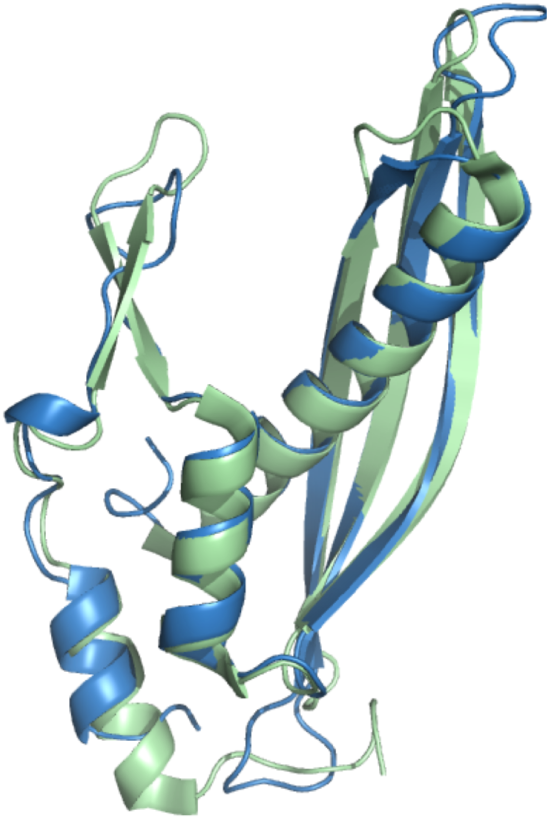
The archaea SSAP A0A2D6XHC3 (green) nearly perfectly aligns with human Rad52 (blue) with a TM-score of 0.82.

## 4 Discussion

### SSAPs across all three superkingdoms

Single-strand annealing proteins form one superfamily and despite deteriorated sequences, there is a common structural motif shared by all of them. Two results stand out in particular: First, the few archaea SSAPs display the same structural diversity as the many bacterial SSAPs. Second, while all the five SSAP families are present in archaea and bacteria, eukaryotes harbor with a few exceptions the Rad52 family only. This is consistent with Williams et al. who posit archaea and bacteria as the two primary domains of life from which eukaryotes arose in partnership [22]. Archaea as eukaryotic ancestry besides bacteria was also investigated by Eme et al., who found that archaea carry several genes formerly believed to be eukaryotic specific [31]. This resonates with our finding of good similarity between eukaryotic and bacterial Rad52, which generally varies in the length of strands in the *β*-sheet motif (see Figure 3) as well as an archaea SSAP, whose *β*-hairpin and *β*-sheet motifs are nearly identical to human. This suggests that SSAP function may be so fundamental that its structural base is heavily constrained across all kingdoms of life. Overall, we were surprised to find only 1% of SSAPs to be archaea. While this is consistent with an estimate of biomass by Bar-On et al. [32], it is inconsistent with Karner et al., who estimate the same order of magnitude of archaea and bacteria cells in the oceans [33]. However, biomass and number of cells may be difficult to relate to the number of sequences for a superkingdom. Therefore, we resolved this issue by counting archaea sequences in the UniProt database. We found 6,606,939 archaea sequences among all 252,170,925 UniProt sequences, which amounts to 2.62%. This means that the 1% archaea SSAPs are only slightly underrepresented in comparison to UniProt in total.

### SSAPs and oligomerization

The advent of large-scale structure prediction has enabled this study, which is the largest structural comparison of the Rad52 superfamily to date. Interestingly, predicted structures not only resemble the overall SSAP structural motif but also aspects of quaternary structure such as the width and general shape of the SSAP monomers, which enables assembly into larger ring or helical structures. However, while experimental structural data suggests these quaternary structures, there is evidence that they only form at very high protein concentrations. Under physiological conditions the rings and helices form only very partially and in a dynamic process tightly linked to DNA binding [34, 35]. Overall, our study supports the possibility of SSAP oligomerization across all of the families and species.

### Defining superfamilies

Our study shows that there is a common structural basis for single-strand annealing, which is only conserved in structure and not in sequence. Generally, our study is an example of how the advent of large-scale structure prediction can enhance the organization of biological knowledge. From experimental data dating back to the 60s, a few single-strand annealing proteins have been directly experimentally identified. The vast majority have been inferred by sequence similarity. It is interesting to quantify this ratio of direct and indirect evidence. E.g. the gene ontology [36] term “DNA double-strand break processing involved in repair via single-strand annealing” has 145 gene products assigned to it (3.1.2024). However, only four of these annotations are inferred from a direct assay (GeneOntology evidence code IDA) and the vast majority are inferred indirectly from “biological aspect of ancestor” (GeneOntology evidence code IBA). Thus the very concept of the SSAP superfamily goes back to a few proteins with direct experimental evidence while the majority of SSAPs are inferred indirectly. This relation of direct and indirect evidence can be obtained from GeneOntology, but it should be made prominent in databases such as InterPro [19]. Another aspect of defining superfamilies concerns the definition of a common structural motif. The *β*-hairpin and *β*-sheet motif have been known from experimental 3D structures [23] and they have been linked to DNA-binding and to function. Thus, from direct experimental observation there was a link from a small part of a protein to its function. This knowledge is crucial for the study of this paper. A large-scale structural search without knowledge of such motifs is difficult. Recently, there has been progress on search of millions of predicted protein structures with the FoldSeek tool [37] and a large-scale clustering of the whole of the AlphaFold database [21], [38]. Yet, the clusterings cannot establish the link between the five SSAP families. One reason that contributes to the difficulty of this is that large portions of the protein outside the motifs vary strongly and can contain large disordered regions that cannot be structurally compared. It is a very fine borderline between unrelated and distantly related. However, this study shows that through the use of sequence, structure, and functional knowledge, it is possible to find this borderline. How to apply this at a large scale to all of the InterPro families is still an open question.

## 5 Conclusion

In conclusion, our study offers a comprehensive examination of the Rad52 single-strand annealing protein (SSAP) superfamily, providing insights into their evolutionary trajectories and structural diversity. By integrating data from the InterPro database and AlphaFold predictions, we deepened our comprehension of SSAP families, including Rad52, Erf, Sak3, Red*β*, and RecT. Noteworthy is our support for the notion that eukaryotic Rad52 likely evolved from predecessors present in both bacterial and archaeal lineages. Furthermore, our analysis highlights the presence of structural similarities among SSAPs, underscoring the significance of conserved motifs in their functionality despite limited sequence conservation. Additionally, the finding of a novel SSAP candidate, an oceanic archaeal SSAP resembling human Rad52, underscores the importance of integrating structural and sequence data in unraveling protein evolution and function. Our findings not only enrich existing knowledge but also unveil new avenues for exploration, providing a robust framework for further investigations into the intricate world of protein evolution and function.

## 6 Methods

Interpro [19] was accessed on 01.08.2023 for families Rad52 (IPR041247), Erf (PF04404), Sak3 (IPR009425), RecT (IPR004590), and Red*β* (IPR010183). AlphaFold (version 3) [20], [21] structures were retrieved on 02.10.2022. Structures with an average pLDDT confidence score lower than 70% were filtered out. Taxonomic information was extracted from UniProt (accessed 01.09.2023). Structures in Figure 1A, 1B, 2, 3, 5, 6, and Supplementary Figure S1A were compared by USAlign’s TM-score [39]. The dendrograms in Figure 1 and in Fig. S1 were created by hierarchical clustering with average linkage using the Python packages skbio and ete3 [40]. They were visualized by iToL (interactive Tree of Life). The four representatives, A0A842UJM1, A0A843GZ30, L0KZL4, A0A2D6XBR4, in Figure 1 were selected in two steps. First, the top four clusters in the hierarchical clustering were selected. Second, for each cluster, the structure, which had the highest average similarity to its cluster members, was selected. For Fig. S1A, sequences were compared using Blast similarity (identity*coverage) using the BLOSUM62 substitution matrix. PyMOL was used for Figure 6 (cealign), Table 3 (secondary structure assignment), and the computation of atom clashes (atoms of one monomer within 2A of atoms of the neighboring monomer). PDB structures 1kn0, 7ub2, and 7ujl served as experimental references.

## 7 Data Availability

All data used to generate this work are accessible via the link provided below. This collection includes both raw and generated data. The raw data consists of PDB files for all SSAP proteins and their FASTA sequences. The generated data encompass BLAST results (sequence data), structural alignments (TM scores), taxonomy information, and family memberships from InterPro. Please refer to the following link to access the data.

Raw data: ./rawData/[PDBs, Fasta]

Alignment results: ./alignments/

https://sharing.biotec.tu-dresden.de/index.php/s/vNrJ3aHLSUN6kZE

## 8 Acknowledgement

We kindly acknowledge financial support from the BMBF projects scads.ai and SNRT as well as access to high-performance computing through the ZIH of TU Dresden.

## 9 Author contributions

AAF, FS and MS conceived the study, AAF and MM implemented the study, AAF, BM, SL, MS analysed data, AAF, FS and MS wrote the manuscript.

## Supplementary Material

**Fig. S1:**
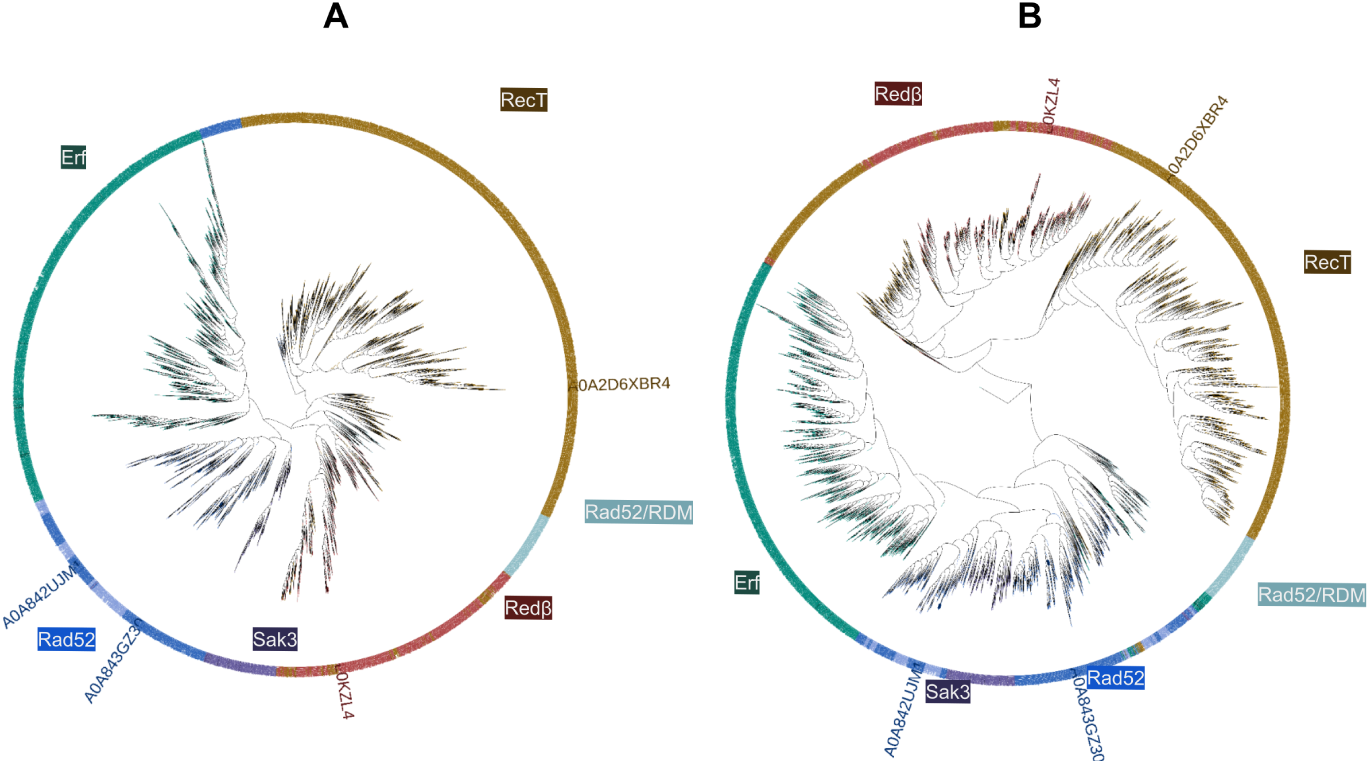
10280 SSAPs clustered by sequence (A) and by structural (B) similarity. Both trees agree by and large.

### Supplementary tables

Tables can be accessed through this link: https://sharing.biotec.tu-dresden.de/index.php/s/vozoYURFI3ctWzP

Table S1: TM-scores for all 10280 SSAPs

Table S2: TM-scores for unreviewed proteins from UniProt with a TM-score of greater 0.7 to at least one SSAP representative.

## References

[1] Mortensen, U.H., Lisby, M., Rothstein, R.: Rad52. Curr Biol 19(16), 676– 677 (2009)

[2] Balboni, B., Rinaldi, F., Previtali, V., Ciamarone, A., Girotto, S., Cavalli, A.: Novel Insights into RAD52’s Structure, Function, and Druggability for Synthetic Lethality and Innovative Anticancer Therapies. Cancers (Basel) 15(6) (2023)

[3] Singleton, M.R., Wentzell, L.M., Liu, Y., West, S.C., Wigley, D.B.: Structure of the single-strand annealing domain of human RAD52 protein. Proc Natl Acad Sci U S A 99(21), 13492–13497 (2002)

[4] Kagawa, W., Kurumizaka, H., Ishitani, R., Fukai, S., Nureki, O., Shibata, T., Yokoyama, S.: Crystal structure of the homologous-pairing domain from the human Rad52 recombinase in the undecameric form. Mol Cell 10(2), 359–371 (2002)

[5] Sugiyama, T., Kantake, N., Wu, Y., Kowalczykowski, S.C.: Rad52-mediated DNA annealing after Rad51-mediated DNA strand exchange promotes second ssDNA capture. EMBO J 25(23), 5539–5548 (2006)

[6] Passy, S.I., Yu, X., Li, Z., Radding, C.M., Egelman, E.H.: Rings and filaments of beta protein from bacteriophage lambda suggest a superfamily of recombination proteins. Proc Natl Acad Sci U S A 96(8), 4279–4284 (1999)

[7] Thresher, R.J., Makhov, A.M., Hall, S.D., Kolodner, R., Griffith, J.D.: Electron microscopic visualization of RecT protein and its complexes with DNA. J Mol Biol 254(3), 364–371 (1995)

[8] Iyer, L.M., Koonin, E.V., Aravind, L.: Classification and evolutionary history of the single-strand annealing proteins, RecT, Redbeta, ERF and RAD52. BMC Genomics 3, 8 (2002)

[9] Erler, A., Wegmann, S., Elie-Caille, C., Bradshaw, C.R., Maresca, M., Seidel, R., Habermann, B., Muller, D.J., Stewart, A.F.: Conformational adaptability of Redbeta during DNA annealing and implications for its structural relationship with Rad52. J Mol Biol 391(3), 586–598 (2009)

[10] Park, M.S., Ludwig, D.L., Stigger, E., Lee, S.H.: Physical interaction between human RAD52 and RPA is required for homologous recombination in mammalian cells. J Biol Chem 271(31), 18996–19000 (1996)

[11] Subramaniam, S., Erler, A., Fu, J., Kranz, A., Tang, J., Gopalswamy, M., Ramakrishnan, S., Keller, A., Grundmeier, G., ller, D., Sattler, M., Stewart, A.F.: is insufficient for homologous recombination and the additional requirements involve intra- and inter-molecular interactions. Sci Rep 6, 34525 (2016)

[12] Lopes, A., Amarir-Bouhram, J., Faure, G., Petit, M.A., Guerois, R.: Detection of novel recombinases in bacteriophage genomes unveils Rad52, Rad51 and Gp2.5 remote homologs. Nucleic Acids Res 38(12), 3952–3962 (2010)

[13] Matsubara, K., Malay, A.D., Curtis, F.A., Sharples, G.J., Heddle, J.G.: . PLoS One 8(11), 78869 (2013)

[14] Newing, T.P., Brewster, J.L., Fitschen, L.J., Bouwer, J.C., Johnston, N.P., Yu, H., Tolun, G.: Red-mediated homologous DNA recombination. Nat Commun 13(1), 5649 (2022)

[15] Caldwell, B.J., Norris, A.S., Karbowski, C.F., Wiegand, A.M., Wysocki, V.H., Bell, C.E.: family recombinase in complex with a duplex intermediate of DNA annealing. Nat Commun 13(1), 7855 (2022)

[16] Al-Fatlawi, A., Schroeder, M., Stewart, A.F.: The Rad52 SSAP superfamily and new insight into homologous recombination. Commun Biol 6(1), 87 (2023)

[17] Illergard, K., Ardell, D.H., Elofsson, A.: Structure is three to ten times more conserved than sequence–a study of structural response in protein cores. Proteins 77(3), 499–508 (2009)

[18] Jumper, J., Evans, R., Pritzel, A., Green, T., Figurnov, M., Ronneberger, O., Tunyasuvunakool, K., Bates, R., dek, A., Potapenko, A., Bridgland, A., Meyer, C., Kohl, S.A.A., Ballard, A.J., Cowie, A., Romera-Paredes, B., Nikolov, S., Jain, R., Adler, J., Back, T., Petersen, S., Reiman, D., Clancy, E., Zielinski, M., Steinegger, M., Pacholska, M., Berghammer, T., Bodenstein, S., Silver, D., Vinyals, O., Senior, A.W., Kavukcuoglu, K., Kohli, P., Hassabis, D.: Highly accurate protein structure prediction with AlphaFold. Nature 596(7873), 583–589 (2021)

[19] Paysan-Lafosse, T., Blum, M., Chuguransky, S., Grego, T., Pinto, B.L., Salazar, G.A., Bileschi, M.L., Bork, P., Bridge, A., Colwell, L., Gough, J., Haft, D.H., Letunic, I., Marchler-Bauer, A., Mi, H., Natale, D.A., Orengo, C.A., Pandurangan, A.P., Rivoire, C., Sigrist, C.J.A., Sillitoe, I., Thanki, N., Thomas, P.D., Tosatto, S.C.E., Wu, C.H., Bateman, A.: Interpro in 2022. Nucleic Acids Res 51(D1), 418–427 (2023)

[20] Jumper, J., Evans, R., Pritzel, A., Green, T., Figurnov, M., Ronneberger, O., Tunyasuvunakool, K., Bates, R., Zidek, A., Potapenko, A., Bridgland, A., Meyer, C., Kohl, S.A.A., Ballard, A.J., Cowie, A., Romera-Paredes, B., Nikolov, S., Jain, R., Adler, J., Back, T., Petersen, S., Reiman, D., Clancy, E., Zielinski, M., Steinegger, M., Pacholska, M., Berghammer, T., Bodenstein, S., Silver, D., Vinyals, O., Senior, A.W., Kavukcuoglu, K., Kohli, P., Hassabis, D.: Highly accurate protein structure prediction with alphafold. Nature 596(7873), 583–589 (2021)

[21] Varadi, M., Anyango, S., Deshpande, M., Nair, S., Natassia, C., Yordanova, G., Yuan, D., Stroe, O., Wood, G., Laydon, A., Zidek, A., Green, T., Tunyasuvunakool, K., Petersen, S., Jumper, J., Clancy, E., Green, R., Vora, A., Lutfi, M., Figurnov, M., Cowie, A., Hobbs, N., Kohli, P., Kleywegt, G., Birney, E., Hassabis, D., Velankar, S.: Alphafold protein structure database: massively expanding the structural coverage of protein-sequence space with high-accuracy models. Nucleic Acids Res 50(D1), 439–444 (2022)

[22] Williams, T.A., Foster, P.G., Cox, C.J., Embley, T.M.: An archaeal origin of eukaryotes supports only two primary domains of life. Nature 504(7479), 231–6 (2013)

[23] Kagawa, W., Kurumizaka, H., Ishitani, R., Fukai, S., Nureki, O., Shibata, T., Yokoyama, S.: Crystal structure of the homologous-pairing domain from the human rad52 recombinase in the undecameric form. Mol Cell 10(2), 359–71 (2002)

[24] Newing, T.P., Brewster, J.L., Fitschen, L.J., Bouwer, J.C., Johnston, N.P., Yu, H., Tolun, G.: Red*β*eta(177) annealase structure reveals details of oligomerization and lambda red-mediated homologous dna recombination. Nat Commun 13(1), 5649 (2022)

[25] Caldwell, B.J., Norris, A.S., Karbowski, C.F., Wiegand, A.M., Wysocki, V.H., Bell, C.E.: Structure of a rect/red*β*eta family recombinase in complex with a duplex intermediate of dna annealing. Nat Commun 13(1), 7855 (2022)

[26] Zhang, Y., Skolnick, J.: Scoring function for automated assessment of protein structure template quality. Proteins 57(4), 702–10 (2004)

[27] Xu, J., Zhang, Y.: How significant is a protein structure similarity with tm-score = 0.5. Bioinformatics 26(7), 889–95 (2010)

[28] Al-Fatlawi, A., Menzel, M., Schroeder, M.: Is protein blast a thing of the past. Nat Commun 14(1), 8195 (2023)

[29] Illergard, K., Ardell, D.H., Elofsson, A.: Structure is three to ten times more conserved than sequence–a study of structural response in protein cores. Proteins 77(3), 499–508 (2009)

[30] Tully, B.J., Graham, E.D., Heidelberg, J.F.: The reconstruction of 2,631 draft metagenome-assembled genomes from the global oceans. Sci Data 5, 170203 (2018)

[31] Eme, L., Spang, A., Lombard, J., Stairs, C.W., Ettema, T.J.G.: Archaea and the origin of eukaryotes. Nat Rev Microbiol 15(12), 711–723 (2017)

[32] Bar-On, Y.M., Phillips, R., Milo, R.: The biomass distribution on earth. Proc Natl Acad Sci U S A 115(25), 6506–6511 (2018)

[33] Karner, M.B., DeLong, E.F., Karl, D.M.: Archaeal dominance in the mesopelagic zone of the pacific ocean. Nature 409(6819), 507–10 (2001)

[34] Kharlamova, M.A., Kushwah, M.S., Jachowski, T.J., Subramaniam, S., Stewart, A.F., Kukura, P., Schäffer, E.: Short oligomers rather than rings of human rad52 promote single-strand annealing. bioRxiv, 2023–08 (2023)

[35] Liang, C.-C., Greenhough, L.A., Masino, L., Maslen, S., Bajrami, I., Tuppi, M., Skehel, M., Taylor, I.A., West, S.C.: Mechanism of single-stranded dna annealing by rad52–rpa complex. Nature (2024)

[36] Aleksander, S.A., Balhoff, J., Carbon, S., Cherry, J.M., Drabkin, H.J., Ebert, D., Feuermann, M., Gaudet, P., Harris, N.L., Hill, D.P., Lee, R., Mi, H., Moxon, S., Mungall, C.J., Muruganugan, A., Mushayahama, T., Sternberg, P.W., Thomas, P.D., Van Auken, K., Ramsey, J., Siegele, D.A., Chisholm, R.L., Fey, P., Aspromonte, M.C., Nugnes, M.V., Quaglia, F., Tosatto, S., Giglio, M., Nadendla, S., Antonazzo, G., Attrill, H., Dos Santos, G., Marygold, S., Strelets, V., Tabone, C.J., Thurmond, J., Zhou, P., Ahmed, S.H., Asanitthong, P., Luna Buitrago, D., Erdol, M.N., Gage, M.C., Ali Kadhum, M., Li, K.Y.C., Long, M., Michalak, A., Pesala, A., Pritazahra, A., Saverimuttu, S.C.C., Su, R., Thurlow, K.E., Lovering, R.C., Logie, C., Oliferenko, S., Blake, J., Christie, K., Corbani, L., Dolan, M.E., Drabkin, H.J., Hill, D.P., Ni, L., Sitnikov, D., Smith, C., Cuzick, A., Seager, J., Cooper, L., Elser, J., Jaiswal, P., Gupta, P., Jaiswal, P., Naithani, S., Lera-Ramirez, M., Rutherford, K., Wood, V., De Pons, J.L., Dwinell, M.R., Hayman, G.T., Kaldunski, M.L., Kwitek, A.E., Laulederkind, S.J.F., Tutaj, M.A., Vedi, M., Wang, S.-J., D’Eustachio, P., Aimo, L., Axelsen, K., Bridge, A., Hyka-Nouspikel, N., Morgat, A., Aleksander, S.A., Cherry, J.M., Engel, S.R., Karra, K., Miyasato, S.R., Nash, R.S., Skrzypek, M.S., Weng, S., Wong, E.D., Bakker, E., Berardini, T.Z., Reiser, L., Auchincloss, A., Axelsen, K., Argoud-Puy, G., Blatter, M.-C., Boutet, E., Breuza, L., Bridge, A., Casals-Casas, C., Coudert, E., Estreicher, A., Livia Famiglietti, M., Feuermann, M., Gos, A., Gruaz-Gumowski, N., Hulo, C., Hyka-Nouspikel, N., Jungo, F., Le Mercier, P., Lieberherr, D., Masson, P., Morgat, A., Pedruzzi, I., Pourcel, L., Poux, S., Rivoire, C., Sundaram, S., Bateman, A., Bowler-Barnett, E., Bye-A-Jee, H., Denny, P., Ignatchenko, A., Ishtiaq, R., Lock, A., Lussi, Y., Magrane, M., Martin, M.J., Orchard, S., Raposo, P., Speretta, E., Tyagi, N., Warner, K., Zaru, R., Diehl, A.D., Lee, R., Chan, J., Diamantakis, S., Raciti, D., Zarowiecki, M., Fisher, M., James-Zorn, C., Ponferrada, V., Zorn, A., Ramachandran, S., Ruzicka, L., Westerfield, M.: The gene ontology knowledgebase in 2023. Genetics 224(1) (2023)

[37] van Kempen, M., Kim, S.S., Tumescheit, C., Mirdita, M., Lee, J., Gilchrist, C.L.M., Soding, J., Steinegger, M.: Fast and accurate protein structure search with foldseek. Nat Biotechnol (2023)

[38] Barrio-Hernandez, I., Yeo, J., Janes, J., Mirdita, M., Gilchrist, C.L.M., Wein, T., Varadi, M., Velankar, S., Beltrao, P., Steinegger, M.: Clustering predicted structures at the scale of the known protein universe. Nature 622(7983), 637–645 (2023)

[39] Zhang, C., Shine, M., Pyle, A.M., Zhang, Y.: Us-align: universal structure alignments of proteins, nucleic acids, and macromolecular complexes. Nat Methods 19(9), 1109–1115 (2022)

[40] Huerta-Cepas, J., Serra, F., Bork, P.: Ete 3: Reconstruction, analysis, and visualization of phylogenomic data. Mol Biol Evol 33(6), 1635–8 (2016)

